# A *Drosophila* Wolfram Syndrome 1 (WFS1) homologue synergises with the intracellular Ca^2+^ release channel, IP_3_R to affect mitochondrial morphology and function

**DOI:** 10.1101/2022.11.10.515972

**Authors:** Rose Sebastian Kunnappallil, Gaiti Hasan

## Abstract

Wolfram syndrome (WFS) is an autosomal recessive neurodegenerative disorder, 90% of which is caused by loss of function of the endoplasmic reticular membrane protein Wolframin or WFS 1. Wolfram syndrome results in Diabetes Insipidus, Diabetes Mellitus, Optic Atrophy, and Deafness (DIDMOAD) in humans. In mammalian cells WFS1 interacts with the ER-localised intracellular Ca^2+^ release channel, Inositol Trisphosphate Receptor 1 (IP_3_R1) required for IP_3_ mediated Ca^2+^ release from the endoplasmic reticulum.

Here, we tested functional interactions between IP_3_R and WFS1 mutants in the context of organismal behaviour and neuronal mitochondrial morphology and physiology in a subset of central dopaminergic neurons of *Drosophila melanogaster*. We show strong genetic interactions between trans-heterozygotes of *wfs1* and *itpr* (IP_3_R) mutants by measuring flight deficits. Over-expression of wild-type cDNAs of either interacting partner, *wfs1*^*+*^ or *itpr*^*+*^ rescued the flight deficits. Cellular studies demonstrate changes in mitochondrial Ca^2+^ entry accompanied by enlarged or swollen mitochondria and decreased mitochondrial content in genotypes that are flight defective. In *wfs1* mutant as well as *wfs1* knockdown conditions a reduction in the number of dopaminergic neurons was observed.

Thus, WFS1 interaction with the IP_3_R is required in flight regulating central dopaminergic neurons of *Drosophila*, for optimal mitochondrial Ca^2+^ entry and maintaining mitochondrial morphology. Our study demonstrates that *Drosophila* can be a good model system to understand the cellular and molecular basis of Wolfram syndrome, its impact on systemic physiology and suggests its use in testing putative pharmaceutical interventions.

## Introduction

Wolfram syndrome (WFS) is a rare autosomal recessive neurodegenerative disorder which affects 1 in 770000 people. WFS was first described by Wolfram and Wagener in 1938 in siblings affected by juvenile-onset diabetes mellitus and optic atrophy. It is also referred to as the DIDMOAD syndrome due to its clinical features of diabetes insipidus (DI), diabetes mellitus (DM), optic atrophy (OA) and deafness (D) (Tranebjaerg et al., 2009). Many patients also develop other symptoms of endocrine deficiencies, ataxia, neurogenic bladder and psychiatric conditions (Cremers et al.,1977). Neurodegeneration of the brain stem ultimately results in death of the patient (Scolding et al., 1996). There are two unrelated proteins that result in the above-mentioned clinical symptoms. Mutations in Wolframin (WFS1) causes the prevalent form of WFS1 (Inoue, 1998), whereas mutations in CISD2 (CDGSH Iron Sulfur Domain 2) causes WFS2 (Amr et al., 2007). Both Wolframin and CISD2 are endoplasmic reticulum (ER) membrane resident proteins and their mutations elicit symptoms of mitochondriopathies (Strom et al.,1998; Rouzier et al.,2017).

The Wolframin protein consist of 890 amino acid residues in humans and 853 in *Drosophila*. It has a central hydrophobic region flanked by two hydrophilic regions at the Carboxy (C) and amino (N) terminus. The functions of WFS1 include responding to endoplasmic reticular stress and promoting ER-Ca^2+^ signalling. ER stress components XBP1 (Kakiuchi et al., 2006) and ATF6β (Odisho et al.,2015) are reported to activate WFS1. In studies with mammalian cells, WFS1 has been shown to interact with the IP_3_R (Inositol triphosphate receptor) (Angebault et al.,2018; Cagalinec et al.,2016). WFS1 in complex with IP_3_R and Neuronal Calcium sensor (NCS1) is thought to help maintain Mitochondria Associated Membrane (MAM) structure (Angebault et al., 2018).

Mutants, knockdowns and expression of a dominant negative version of the *Drosophila* IP_3_R all show impaired flight (Banerjee et al., 2004; Sharma and Hasan, 2020). Previous studies have identified a requirement for the IP_3_R in a subset of central Dopaminergic (DA) neurons marked by TH D’GAL4 (Ravi et al., 2018) for maintenance of longer flight bouts (Ravi et al., 2018; Sharma and Hasan, 2020). DA neurons are also affected by a deficiency of WFS1 in mice, in that they show reduced dopamine output and dopamine transporter expression (Matto et al., 2011, Visnapuu et al., 2013). Interaction between WFS1 and D1-like dopamine receptor signalling has also been shown (Tekko et al., 2017).

Since the symptoms of WFS1 mutations are similar to those of mitochondriopathies, several studies have investigated mitochondrial function. Disrupted mitochondrial transportation, fission-fusion cycles, and more frequent mitophagy are observed in WFS1 deficient conditions in mammals (Cagalinec et al., 2016). Mitochondrial abnormalities are also a hallmark of various other neurodegenerative diseases including Parkinson’s, Amylotrophic lateral Schelrosis, Alzheimer’s, Ataxia Telangiactasia and Huntington’s (Lezi and Swerdlow, 2012).

In *Drosophila*, neuronal knockdown of the *wfs1* homologue *(CG 4917)* induces neurodegeneration and behavioural deficits whereas *wfs1* mutants exhibit increase in susceptibility to oxidative stress, axon degeneration and early death. These phenotypes are exacerbated by additional knockdown of *wfs1* in glia (Sakakibara et al., 2018). As WFS1 in mammalian cells interacts with the IP_3_R, we hypothesised that *Drosophila* could be a good model system to further investigate the systemic and cellular effects of IP_3_R/WFS1 interactions. We establish that WFS1 and IP_3_R are required for Ca^2+^ entry into mitochondria of central dopaminergic neurons as well as normal mitochondrial morphology. The disruption of these cellular functions in DA neurons affects motor function in an age-dependent manner.

## Materials and Methods

### Drosophila stocks used

The *wfs1*^*e03461*^ mutants (BDSC 18157), *wfs1*^*MI14041*^ mutants (BDSC 59250), *UAS wfs1* (BDSC 8357, BDSC 8356), *wfs1* RNAi (BDSC 53330), *UAS JRCaMP* (BDSC 63793) and nSybGal4 (BDSC 51635) were obtained from Bloomington Drosophila Stock centre. *itpr RNAi* (*UASdsitpr* (1063R-2) line) was obtained from from the National Institute of Genetics Fly Stock Center, Japan. The *itpr* mutants used for the study *itpr*^*KA1091*^, *itpr*^*UG3*^, *itpr*^*KA901*^ and *itpr*^*SV35*^ were ethyl methanesulfonate (EMS) mutants generated in the lab (Joshi *et al*.,2004). Mito-GCaMP flies were provided by Prof. Fumiko Kawasaki, *TH GAL4* (Friggi-Grelin et al., 2003) by Serge Birman (CNRS, ESPCI Paris Tech, 594 France), *THD’* GAL4 (Liu et al., 2012) from Mark N Wu (Johns Hopkins University, Baltimore). The fly stocks used were grown in corn meal agar medium at 25°C and used for the experiments.

### Flight assay

To analyse the effect of the deficiency created by RNAi or mutation on flight, we used the flight assay protocol first described by Manjila and Hasan (2019). Flies of the required genotype are grown along with appropriate controls. For knockdown experiments with RNAi the controls included GAL4 lines and UAS lines crossed to Canton-S whereas for mutant studies we tested heterozygotes of individual mutants as controls. Freshly eclosed flies are collected and aged for 5, 10 or 20 days as per experiment. These flies are anesthetised on ice for 2 min and tethered onto a thin metal wire using nail polish. The experiment is done in batches of 10 flies at a time. After recovery from cold anaesthesia, a single gentle air puff is given and the duration of a continuous flight bout is recorded for a maximum of 15 minutes. A minimum of 30 flies are used per genotype. The data so generated was tabulated as flight duration in seconds and plotted as box plots using Origin (OriginLab, Northampton, MA). Statistical analysis (Mann-Whitney-U test) was also performed using Origin.

### Calcium imaging

Whole brains from 5-day old flies expressing genetic Ca^2+^ reporters for cytosolic Ca^2+^ (*UAS JRCaMP*) and mitochondrial Ca^2+^ (UAS *Mito-GCaMP*) in appropriate genetic backgrounds were dissected in AHL (Adult Haemolymph-Like) solution (108 mM NaCl, 5 mM KCl, 2 mM CaCl_2_, 8.2 mM MgCl_2_, 4 mM NaHCO_3_, 1 mM NaH_2_PO_4_, 5 mM trehalose, 10 mM sucrose, 5 mM Tris, pH 7.5). The dissected brains were mounted in 0.8% low melt agarose placed in a small hole at the bottom of a 35mm Petri dish with a coverslip attached below the hole for facilitating microscopic analysis. To the agarose embedded brains 100 µl of AHL buffer was added. Dopaminergic neurons in the brains were subjected to time-lapse imaging at an interval of 2 seconds (Olympus FV3000 confocal microscope, Olympus Corp) using 40X magnification. Brains were stimulated with AHL and 50µM Carbachol added at the 50^th^ sec and 100^th^ sec respectively. f. Both the reporters were driven by *TH D’ GAL4*. The time-lapse images so obtained were processed using Image-J software. The ROIs containing each cell were measured for fluorescence intensities. The ratio of change in fluorescence to baseline fluorescence (ΔF/F) were measured and plotted as a graph across time frames using Origin software.

### Immunohistochemistry

Adult fly brains were dissected in PBS (Phosphate buffered Saline; pH7.2), fixed in 4% PFA (Paraformaldehyde) and washed in 0.3% PBST (PBS + 0.3% Triton-X-100) 3 times. This was followed by blocking using NGS (Normal Goat Serum) or FBS (Fetal Bovine Serum) for 30 minutes, incubation in primary antibody (overnight, 4°C, anti-TH-1:200 and anti-RFP 1:500) and secondary incubation (2 hours, at room temperature, 1:500). Each of the above steps were followed by 3 washes in 0.3% PBST. The sample was then mounted in 60% glycerol and imaged on Olympus FV3000 confocal microscope (60X, Zoom-3X). The images were obtained as Z-stacks of 0.5 µm thickness each. The sources of antibodies are TH (Immunostar), anti-RFP, Alexa Fluor-488 and Alexa Flour-594 (Life Technologies).

### Analysis of mitochondrial morphology

The images were processed using image-J software. Maximum intensity projections obtained from Z-stacks were used for counting TH positive cells and to analyse mitochondrial morphology in PPL1 cluster of neurons. The cells with mitochondrial DsRed expression as well as swollen mitochondria were counted for each genotype. The data was statistically analysed and plotted using Origin.

### Statistical analysis

For non-parametric data, the p-values were determined using Mann-Whitney U test (pairwise comparisons). Normal distributions were subjected to Student’s t-test. For comparing multiple genotypes Kruskal -Wallis test was used. The statistical analysis was performed by using Origin software.

### Data availability

All data presented in manuscript can be found in associated files uploaded.

## Results

### WFS1 is required for flight in *Drosophila melanogaster*

To test whether WFS1 is required for maintaining neuronal health in *Drosophila* we tested initiation and maintenance of flight in homozygous mutants of *wfs1*. Previous studies from the lab had indicated a neuronal requirement of IP_3_R for flight (Banerjee et al., 2004). Since mammalian studies have shown that WFS1 is an interacting partner of the IP_3_R1 (Angebault et al., 2018) we tested *wfs1* deficiency flies for flight, a phenotype described previously for IP_3_R mutants, with a focus in dopaminergic neurons (Pathak et al., 2015 and Sharma and Hasan, 2020).

The homozygous null mutant of *wfs1* (*wfs1*^*e03461*^/ *wfs1*^*e03461*^) was flightless (Video 1, Fig 1A and 1B). We also used a hypomorphic C terminal truncated mutant of *wfs1* (*wfs1*^*MI14041*^) in which the flight defect manifested after ageing for 20 days (Fig 1C). Interestingly, in human patients, mutations towards the C-terminal tend to be milder and result in later age of onset, whereas mutations towards the N-terminal are lethal at earlier age and present severe symptoms (Matsunaga et al., 2014, Rohayem et al., 2011). Heteroallelic combination of *wfs1*^*e03461*^/ *wfs1*^*MI14041*^ resulted in flight defects as well (Supp 1A) i dismissing non-specific effects of the *wfs1* mutant alleles tested here. To test if the flight deficit in *wfs1* mutants had a neuronal focus we generated flies with pan-neuronal *nSybGAL4* driven knockdown of *wfs1*. A strong flight deficit with a mean flight time of 200 secs was observed in flies with pan-neuronal knockdown of *wfs1* as compared to the control strains where mean flight times were closer to 600 secs (Fig 1D). The pan-neuronal knockdown of *wfs1* resulted in flight defects starting at 5 days. Further we tested the effect of *wfs1* knock down in a majority of dopaminergic neurons (*TH GAL4;* Friggi-Grelin et al., 2003) and in a previously identified dopaminergic subset (*THD’ GAL4*; Liu et al., 2012) where the IP_3_R is required for maintenance of long flight bouts (Sharma and Hasan, 2020). RNAi mediated knockdown of *wfs1* by either *THGAL4* or *THD’GAL4* exhibits a flight defect comparable to *itpr RNAi* at age 5 days and 10 days (Fig 1D). Though, pan-neuronal knockdown of *wfs1* gave similar flight deficits knockdown in dopaminergic neurons at 5 and 10 days by 20^th^ day there was a marked improvement in maintenance of flight bouts in flies with *wfs1 RNAi* driven by *TH GAL4* and *TH D’ GAL4* on the 20^th^ day (Supp 1B). The use of glutamatergic *OK371 GAL4* also resulted in flight defects (Supp 1C).

**Figure 1:**
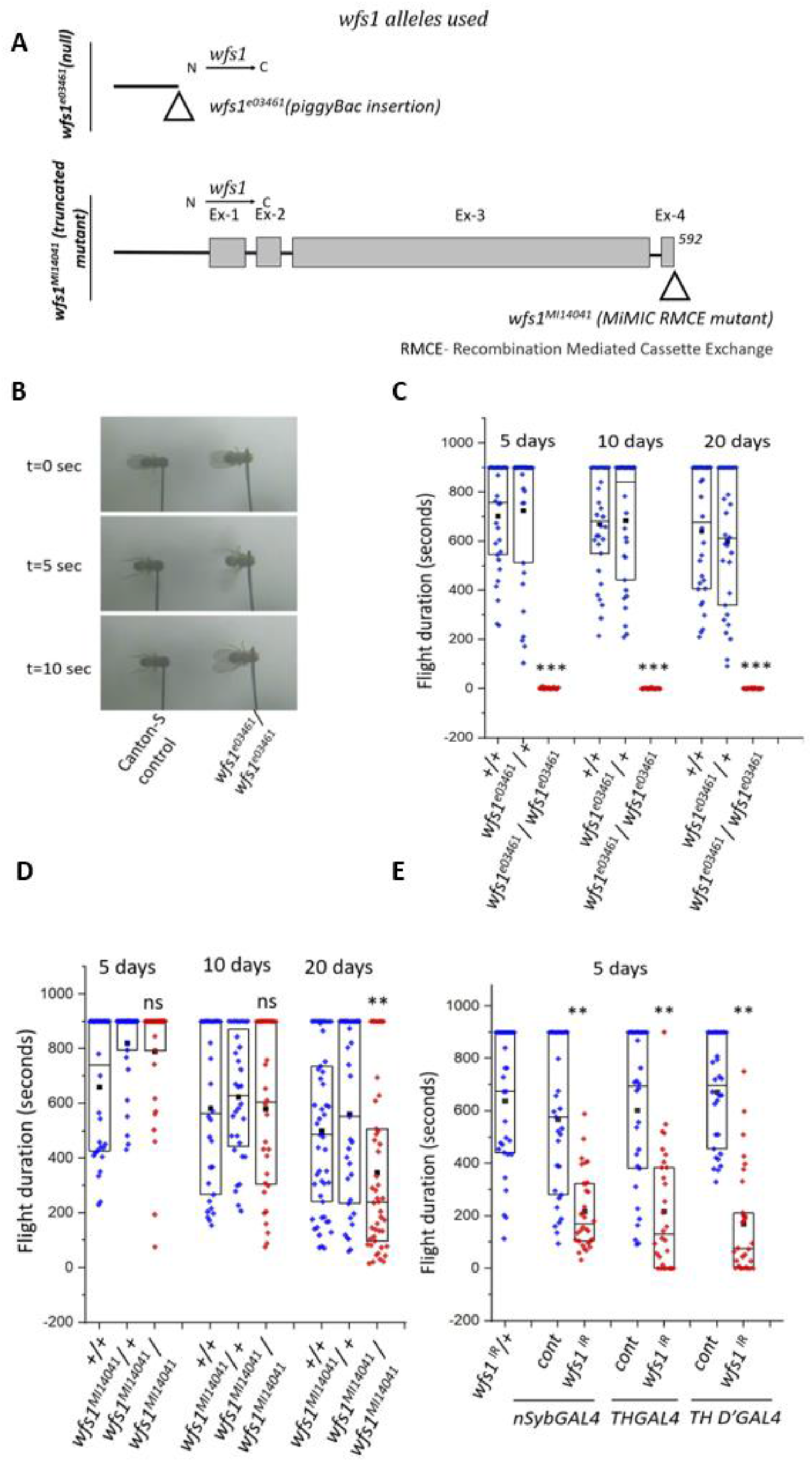
WFS1 is required for flight in dopaminergic neurons of *Drosophila melanogaster*. (A) Graphical representations of a *wfs1* null mutant *wfs1*^*e03461*^ and a truncated mutant (*wfs1*^*MI14041*^)(B) *wfs1* null mutants show a loss of flight (C) Graphical representation of flight duration in a *wfs1* null mutant and (D) a 592 aa C-terminal truncated mutant of *wfs1* (E) *GAL4 –UAS RNAi* mediated knockdown of *wfs1* using either pan neuronal *GAL4 (nSyb)*, or dopaminergic GAL4s, *THGAL4* and *TH D’GAL4* (5-days old flies) (individual data points represent flight durations of each fly, black dot and horizontal line represent mean and median respectively and box size represents range of 25-75% (N ≥30, *** p-values < 0.001, ** p-values 0.001-0.005, Mann-Whitney test).

### WFS1 interacts with IP_3_R as shown by transheterozygotes of WFS1 and IP_3_R mutants

To understand if WFS1 and IP_3_R function together in neurons, we performed genetic interaction studies. For this, we used transheterozygotes of *wfs1* and *itpr* mutants. The *itpr* alleles used were ethyl methanesulfonate (EMS) mutants generated previously in the lab. The homozygotes of *itpr* ^*ka901*^ and *itpr*^*sv35*^ are lethal (Joshi *et al*.,2004). Trans-heterozygotes of the *wfs1* null mutant and an *itpr* allele with a point mutation in the Ca^2+^ channel *(wfs1*^*e03461*^/+; *itpr*^*ka901*^*/+*) were tested for flight bout durations (Fig 2A). The controls, heterozygotic *wfs1*^*e0346*1^/+ and *itpr*^*ka901*^*/+* exhibit normal flight durations. The trans-heterozygotes with single copies of each mutant allele exhibit flight defects indicating genetic interaction between the *wfs1* and *itpr* (Fig 2A). Interestingly, there was an age dependent decrease in flight durations from 5^th^ day (417 ± 53.23 s) to 20^th^ day (71.23 ± 20.96 s)

**Figure 2:**
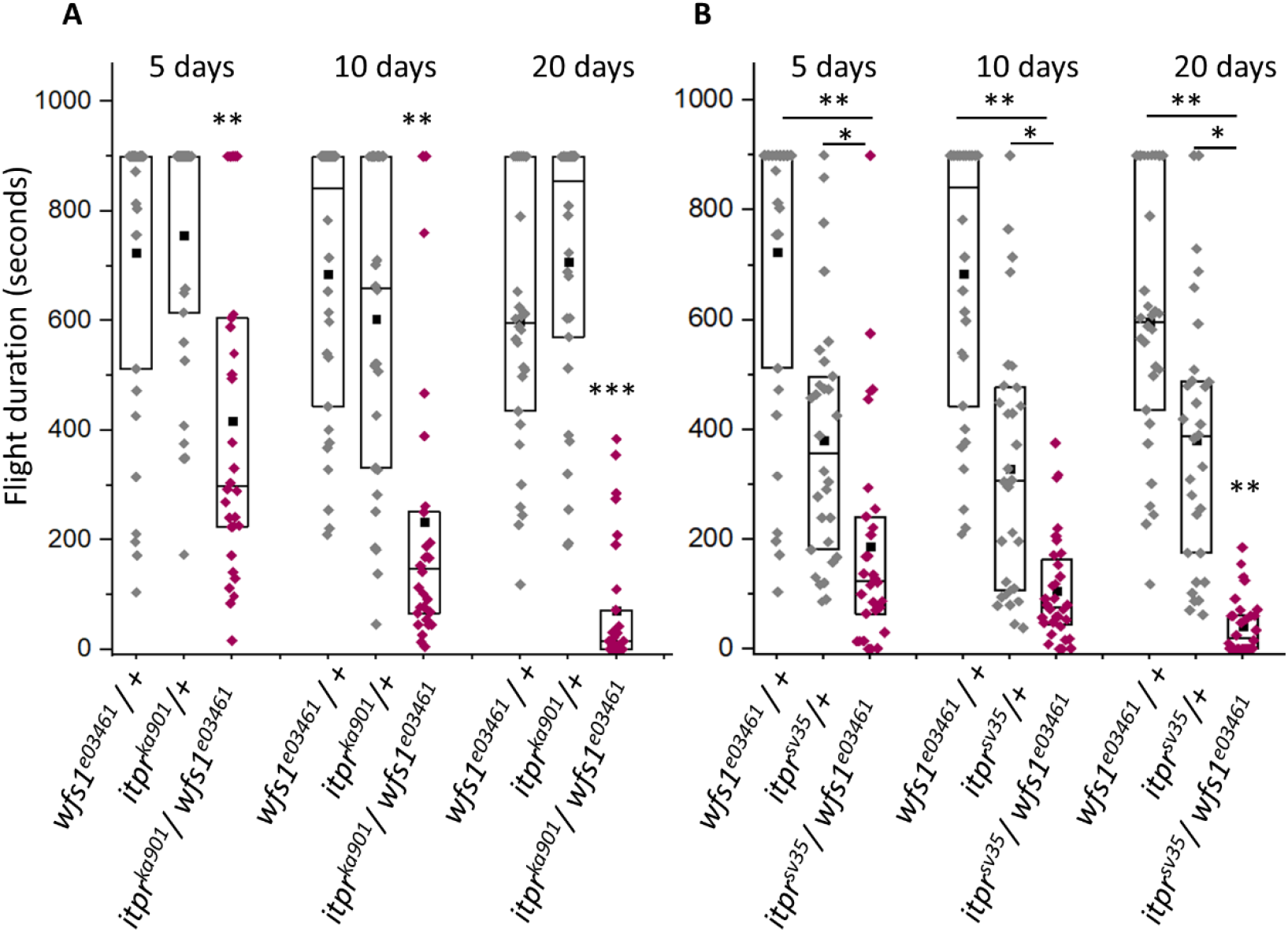
Age dependent loss of flight in transheterozygotes of WFS1 and IP_3_R mutants. Graphical representation of flight duration in transheterozygotes of *wfs1* null mutants in combination with (A) *itpr* ^*KA901*^ harbouring a point mutation in the Ca^2+^ channel domain and (B) *itpr* ^*SV35*^, a truncated null mutant of IP_3_R containing 1572 aa (N ≥30, * p-values < 0.05, ** p-values 0.001-0.005, Mann-Whitney test).

Next, we corroborated the genetic interaction between *wfs1* and *itpr* by using a different mutant, *itpr*^*sv35*^. This allele of *itpr* harbours a stop codon in the modulatory domain of the IP_3_R. This results in a truncated IP_3_R subunit of 1522 amino acids and is a functional null (Joshi et al., 2004). *itpr* ^*sv35*^ in combination with the *wfs1* null (*wfs1*^*e0346*1^/+; *itpr* ^*sv35*^/+) gave an exacerbated flight defect at 5 days (mean flight time with SE) that became worse by the 20^th^ day (mean flight time with SE) in comparison with either heterozygote (Fig 2B). *itpr* ^*sv35*^/+ heterozygotes also exhibit flight defects when tested for long flight bouts but these were relatively mild compared to the deficit observed in the transheterozygotes (*wfs1*^*e0346*1^/+; *itpr* ^*sv35*^/+). These have not been reported earlier..

Further, a hypomorphic allele of wfs1 (*wfs1* ^*MI14041*^, truncated protein of 592 amino acids) was used in combination with the *itpr* ^*ka901*^ and *itpr* ^*sv35*^. These heteroallelic combinations of *(wfs1*^*MI14041*^/+; *itpr*^*ka901*^*/+* and *wfs1* ^*MI14041*^/+; *itpr* ^*sv35*^/+) also exhibit enhanced flight defects (Supp 2) as compared to corresponding heterozygotes. These data further confirm a genetic interaction between *wfs1* and *itpr* and suggest that the two proteins could function in the same cellular pathway.

### *TH D’ GAL4* mediated overexpression of IP_3_R and WFS1 rescues the flight defect caused by their mutant trans heterozygotes and by knockdown of either gene

Previous reports have suggested that IP_3_R expression in *wfs1* deficient conditions restores Ca^2+^ response, development and axonal growth in primary cortical neurons of rat (Cagalinec *et al*., 2016). In *Drosophila UAS itpr* ^*+*^ overexpression in a dopaminergic neuron subset is sufficient to rescue flight defects in the flightless heteroallelic *itpr* ^*ka1091/ug3*^(*itpr*^*ku*^) mutant combination (Sharma and Hasan, 2020). Therefore we tested the effect of *TH D’ GAL4* (Liu *et al*., 2012) driven overexpression of wildtype cDNA constructs of either gene (*UAS itpr*^*+*^ and *UAS wfs1*^*+*^) on flight deficits in the mutants and knockdowns.

Overexpression of either *UAS itpr*^*+*^ or *UAS wfs1*^+^ in flight defective *wfs1*^*e03461*^/+; *itpr*^*ka901*^/+ trans heterozygotes partially rescued and restored the flight duration (Fig 3 A and Supp 3A). The overexpression of *UAS itpr*^*+*^ in flightless *wfs1* null mutants also restored flight time, but not to the extent of wildtype controls. However, the overexpression of *UAS wfs1*^+^ in flightless *itpr*^*ku*^ (*itpr* ^*ka1091/ug3*^) mutant flies did not result in any rescue (Supp 3B) indicating that the IP_3_R may function downstream to WFS1.

**Figure 3:**
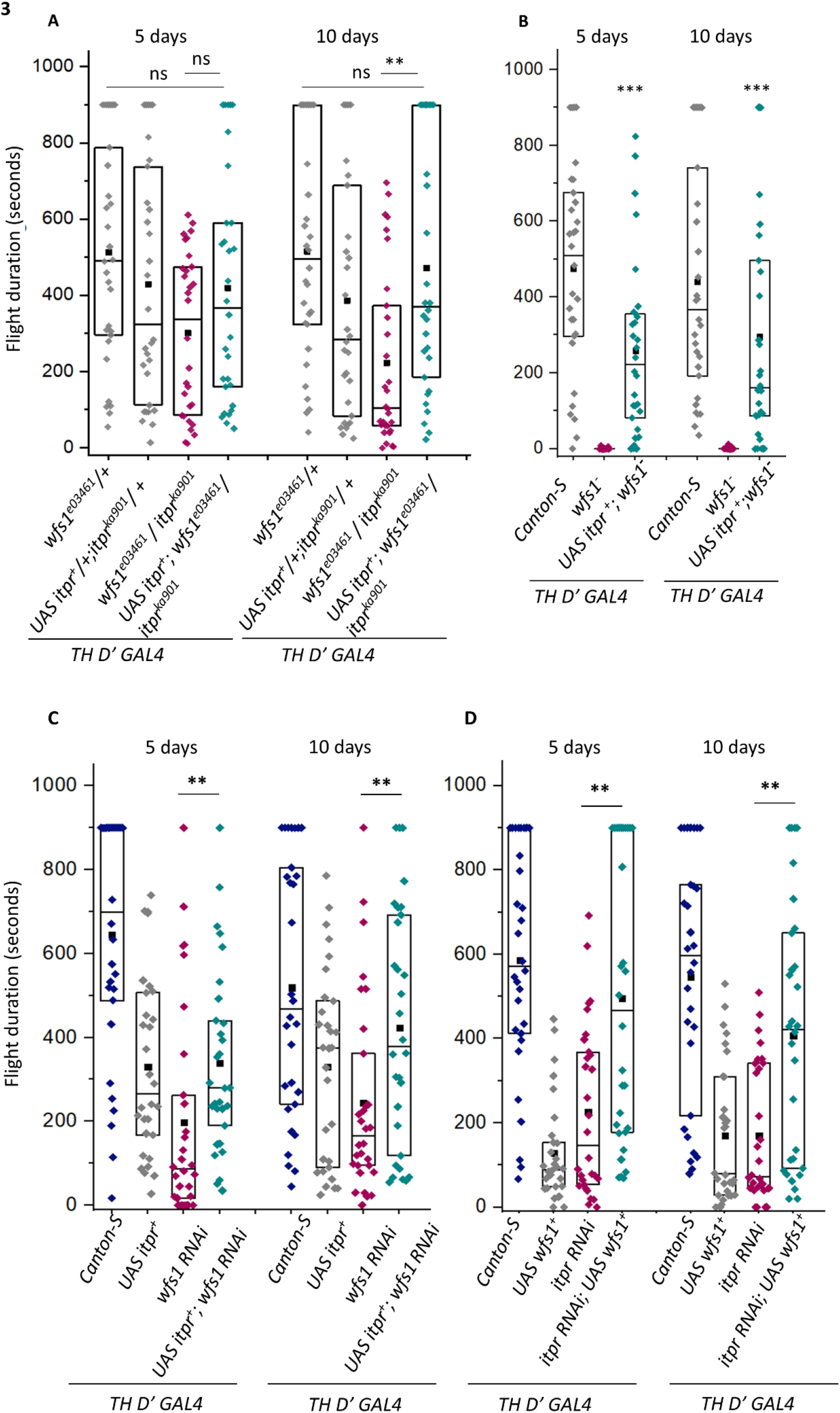
*TH D’ GAL4* mediated overexpression of IP_3_R and WFS1 rescues the flight defect caused by either gene deficiency and of their mutant transheterozygotes. Rescue of (A) *wfs1*^*e03461*^/+; *itpr*^*ka901*^/+ by overexpression of IP_3_R in TH D’ neurons (B) flightless null mutants of *wfs1* (*wfs1*^*e03461*^/ *wfs1*^*e03461*^) by *TH D’* mediated overexpression of *UAS itpr* and (C) *wfs1* knockdown by overexpression of *UAS itpr*. (D) Rescue of *itpr* knockdown by overexpression of *UAS wfs1*. (N ≥30, *** p-values < 0.005, ** p-values 0.001-0.005, Mann-Whitney test)

To further validate the specificity of deficits in dopaminergic neurons, we expressed *UAS itpr*^*+*^ in *wfs1 RNAi* and *UAS wfs1*^*+*^ in *itpr RNAi* backgrounds under the control of *TH D’GAL4*. These experiments resulted in improved flight in both rescue conditions when compared to the knockdown condition and over-expression alone (Fig 3C and 3D). Moreover, *TH D’* mediated overexpression of either *UAS itpr*^*+*^ or *UAS wfs1*^*+*^ resulted in significant flight defects indicating that excessive expression of these proteins can interfere with the normal function of TH D’ dopaminergic neurons.

From these findings we conclude that the action of WFS1 is required for normal function of IP_3_R. However, WFS1 function requires a minimum level of wild type IP_3_R which is very likely present in the *itpr* knock down condition (Fig 3D) but absent in the *itpr*^*ku*^ condition (Supp Fig 3B). These data indicate that IP_3_R function is downstream of WFS1 in dopaminergic neurons of Drosophila similar to a previous finding in rat primary neuronal cell cultures and mouse cortical neurons (Osman *et al*., 2003, Cagalinec *et al*., 2016).

### Endoplasmic Reticular Ca^2+^ release through the IP_3_R into the cytoplasm as well as Ca^2+^ entry in the mitochondria are affected in WFS1 deficient neurons

To understand the cellular basis of flight deficits observed in WFS1 deficient flies, we studied Ca^2+^ responses in mitochondria and cytoplasm of PPL1 dopaminergic neurons with knockdown of *wfs1*. PPL1 neurons are clusters of 12 pairs of neurons each present in the *Drosophila* brain (Mao, 2009). They are marked by *THD’GAL4* (Liu et al., 2012). These neurons require IP_3_R function for maintenance of long flight bouts as described earlier (Pathak et al., 2015; Sharma and Hasan, 2020).

A genetically encoded cytosolic calcium sensor, *UAS JRCaMP6* and a mitochondrial Ca^2+^ sensor, UAS *MitoGCaMP* were used to measure ER-Ca^2+^ release and mitochondrial Ca^2+^ uptake in *TH D’ GAL4* marked PPL1 dopaminergic neurons. The Ca^2+^ measurements were made in ex-vivo brain preparations from wildtype, *wfs1* knockdown and *itpr* knockdown conditions.

Cytosolic and mitochondrial Ca^2+^ transients were measured upon stimulation of PPL1 neurons by carbachol (carbamylcholine), a muscarinic acid receptor (mAChR) agonist (Pathak et al., 2015; Sharma and Hasan, 2020). Addition of carbachol results in a cascade of processes where carbachol binds to its receptor mAChR which in turn activates PLC-β and ultimately leads to increased production of IP_3_ ligands (Millar *et al*., 1995; Srikanth *et al*., 2006), followed by IP_3_R specific Ca^2+^ release from the ER.

Simultaneous measurement of cytosolic Ca^2+^ and mitochondrial Ca^2+^ in response to carbachol (measure of ER-Ca^2+^ release) indicated a decrease in both Ca^2+^ responses in *wfs1 RNAi* and *itpr RNAi* conditions (Fig 4A, 4G). To further understand the Ca^2+^ transients in wildtype versus knockdown conditions, we counted cells that gave a Ca^2+^ response (responders) versus cells that gave no response upon addition of carbachol (non-responders) in each genotype tested. Fewer cells responded in the knockdown conditions.. The percentage of carbachol-induced *JRCaMP* responders (cytosolic Ca^2+^) went down from 42% in wildtype to 30% in *wfs1* RNAi and 32 % in *itpr* RNAi (Fig 4B). *MitoGCaMP* responders, measured simultaneously, went down from 70 % in wildtype to 22 % in *wfs1* RNAi and 50 % in *itpr* RNAi (Fig 4 H). Responders were separated into cells where Ca^2+^ transients were early (< 10s), intermediate (10-20s) and late (>20s) after carbachol addition. Both the cytosolic (*JRCaMP*) response and mitochondrial (*MitoGCaMP*) responses exhibit slower rate of increase in fluorescence (ΔF/F) in *itpr RNAi* and *wfs1 RNAi* in the early responders (Fig 4 C, I). Intermediate responders were not observed consistently in *wfs1* and *itpr* knockdown conditions (Fig 4D, 4J.) Ca^2+^ responses in late responders (Fig 4E, 4K) and fluorescent curves of non-responders (Fig 4F, 4L) appeared similar between wildtype and knockdown conditions. Moreover, the kinetics of Ca^2+^ transients in knockdown conditions were different as compared to wildtype in the responders. Interestingly, after the initial delay, the few responders in *wfs1 RNAi* exhibit a consistently high ΔF/F whereas, the wildtype cells gave a gradual decrease in fluorescence suggesting unregulated ER Ca^2+^ release and mitochondrial Ca^2+^ uptake in *wfs1* knockdown condition (Fig 4C and 4I). The results indicate impairment of both cytosolic and mitochondrial Ca^2+^ response in *wfs1 RNAi* with a greater effect on the mitochondrial response in terms of the number of responders. Similarly, *itpr RNAi* cells respond later compared to wildtype and *wfs1 RNAi*. The difference is more pronounced in Mitochondrial *MitoGCaMP* (Fig 4I and 4J). However, higher proportion of *itpr RNAi* cells responded earlier in case of cytosolic *JRCaMP*(Fig 4C and 4D).

**Figure 4:**
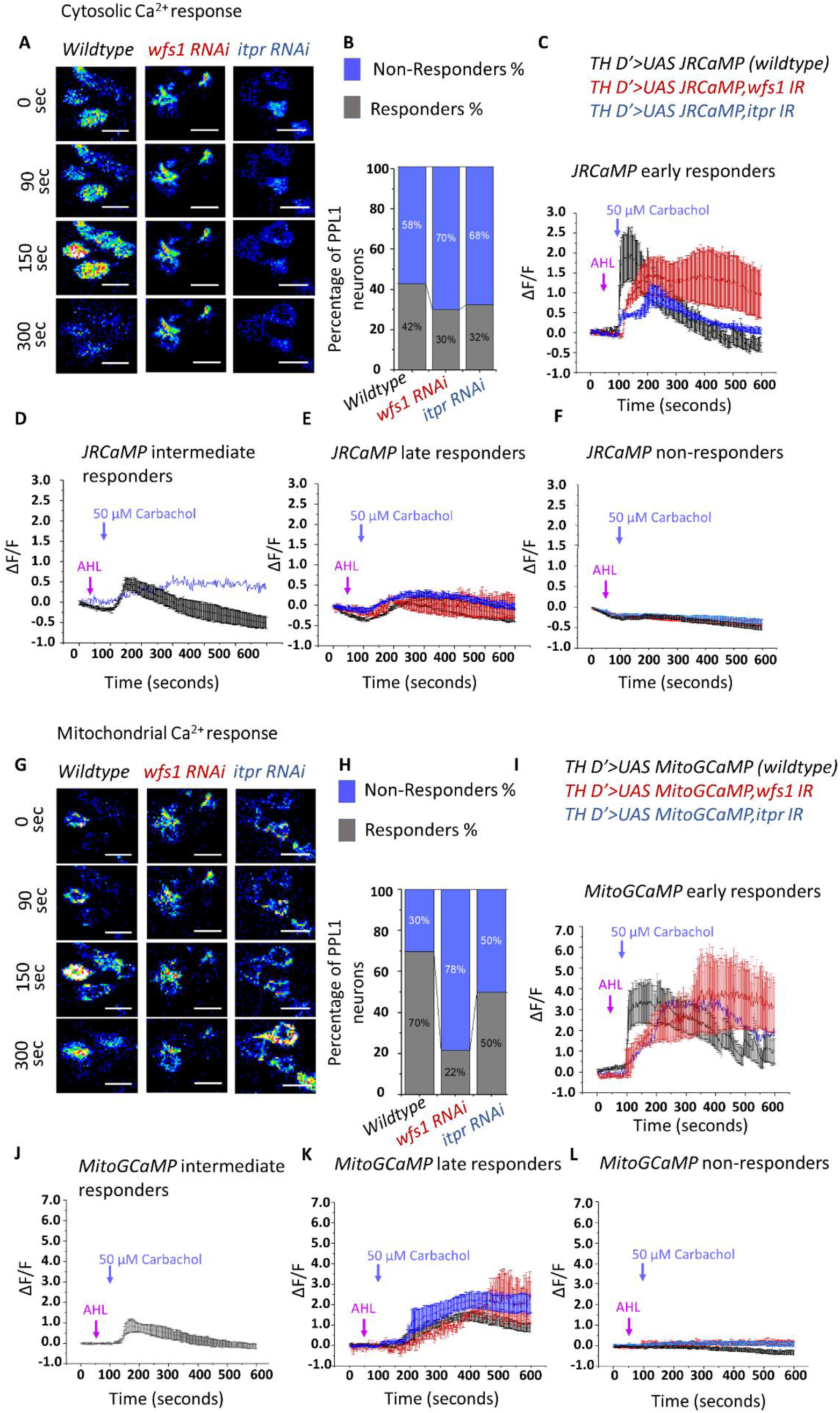
Endoplamic Reticular Ca^2+^ release through the IP_3_R in the cytoplasm and mitochondrial Ca^2+^ entry are affected in WFS1 deficient central dopaminergic neurons. (A-F) Representative images and quantification of cytoplasmic Ca^2+^ transients observed by time-lapse imaging of PPL1 neurons expressing the genetically encoded Ca^2+^ indicator *JRCaMP (THD’ GAL4>UAS JRCaMP)* in response to carbachol. (B) Histogram with percentage of responder and non-responder neurons in the PPL1 cluster of wildtype (n=31), *wfs1* RNAi (n=36) and *itpr* RNAi (n=29) flies. (C-F) Graphical representation of changes in *JRCaMP* fluorescence with time upon addition of Carbachol. (C) early responders, within 10 sec of carbachol addition. Wildtype (n=6), wfs1 RNAi (n=7) and itpr RNAi (n=5). (D) intermediate responders within 10-20 sec after Carbachol addition. Wildtype (n=4), wfs1 RNAi (n=0) and itpr RNAi (n=1). (E) late responders, after 20 sec. Wildtype (n=2), wfs1 RNAi (n=3) and itpr RNAi (n=4) and (F) non-responders. Wildtype (n=19), wfs1 RNAi (n=26) and itpr RNAi (n=19). (G-L) Representative images and quantification of mitochondrial Ca^2+^ transients in PPL1 neurons of the indicated genotypes observed by time-lapse imaging of the mitochondrial Ca^2+^ indicator *MitoGCaMP (THD’ GAL4>UAS MitoGCaMP*) in response to carbachol. (G) representative images of *MitoGCaMP* responses in the indicated genotypes. (H) Histogram with percentage of responder and non-responder neurons in the PPL1 cluster of wildtype (n=31 cells), *wfs1* RNAi (n=36 cells) and *itpr* RNAi (n=29 cells) flies. (I-L) Graphical representation of changes in *MitoGCaMP* fluorescence with time upon addition of Carbachol. (I) Early responders, within 10 sec. Wildtype (n=9), *wfs1* RNAi (n=5) and *itpr* RNAi (n=1). (J) intermediate responders, 10-20 sec. Wildtype (n=7), *wfs1* RNAi (n=0) and *itpr* RNAi (n=0). (K) late responders, after 20 sec. Wildtype (n=5), *wfs*1 RNAi (n=2) and *itpr* RNAi (n=10). (F) Non-responders. Wildtype (n=10), *wfs1* RNAi (n=29) and *itpr* RNAi (n=18). n = number of cells in each genotype.

### Mitochondrial defects and decrease in dopaminergic neurons are found in *wfs1* deficient condition

The disruption of Ca^2+^ homeostasis can affect turnover dynamics of mitochondria and in turn result in degeneration of the cells as exemplified by various neurodegenerative diseases (Rangaraju et al., 2019),while changes in cytosolic Ca^2+^ affect the fission - fusion dynamics of mitochondria (Hom et al., 2010). Mitochondrial Ca^2+^ is a co-factor for mitochondrial enzymes (Rossi *et al*., 2019) and also modulates mitophagy (Barazzuol et al., 2020). Moreover, mitochondria function as asink for Ca^2+^ released from the Endoplasmic reticulum. Altered Ca^2+^ dynamics observed in *wfs1* and i*tpr* mutant alleles and knockdowns could thus affect mitochondrial morphology. To investigate mitochondrial morphology of neurons in the PPL1 cluster we expressed *UAS MitoDsRed* driven by *THD’ GAL4* and immunostained for Tyrosine hydroxylase (TH) to mark dopaminergic neurons. The DsRed channel was converted to 8-bit images and thresholded such that mitochondrial morphology is visible. Healthy mitochondria appear rod-shaped or punctate, whereas, mitochondria that are unhealthy or undergoing mitophagy appear round and enlarged (Valente et al., 2017).

In wildtype PPL1 neurons the mitochondria appear finely grained with thread like connections (Fig 5A). *wfs1* deficient genotypes that showed strong flight defects were investigated for possible mitochondrial abnormalities. *itpr*^*ka901*^ /*wfs1*^*e03461*^ trans heterozygotes and *wfs1*^*e03461*^*/wfs1*^*e0346 1*^homozygous *wfs1* null combinations (both exhibit a marked decrease in the number of mitochondria within THD’ cells (Figure 5B and 5C). The reduction in mitochondria was quantified by measuring *TH D’*>*MitoDsred* expression. In the PPL1 cluster of neurons *MitoDsred* expression among the TH positive cells reduced from an average of 7.6+/-0.41 in wildtype cells to 5.67 +/-0.53 in *wfs1*^*e03461*^*/*+ heterozygotes and 6.67 in *itpr*^*ka901*^ /+ to 3.5 in *wfs1*^*e03461*^*/wfs1*^*e03461*^ (Fig 5F).The lack of 100% *MitoDsRed* expression in control cells could be due to ineffective expression of the *UAS MitoDsRed* construct in the *TH D’* subset of neurons. Moreover the few mitochondria that were present in *wfs1*^*e03461*^*/wfs1*^*e03461*^ had a swollen morphology as represented in Fig 4G. In addition, the number of TH positive cells per PPL1 cluster went down in *wfs1* deficient brains (5.72+/-0.92 in *wfs1*^*RNAi*^ and 6.8+/-0.77 in_*wfs1*^*e03461*^ *mutants* suggesting an effect of mitochondrial function on Tyrosine Hydroxylase (TH) expression (Fig 5F and 5 G). In *itpr*^*ka901*^*/wfs1*^*e03461*^ heterozygotes however the number of TH positive cells were statistically unchanged (Fig 5F).

**Figure 5:**
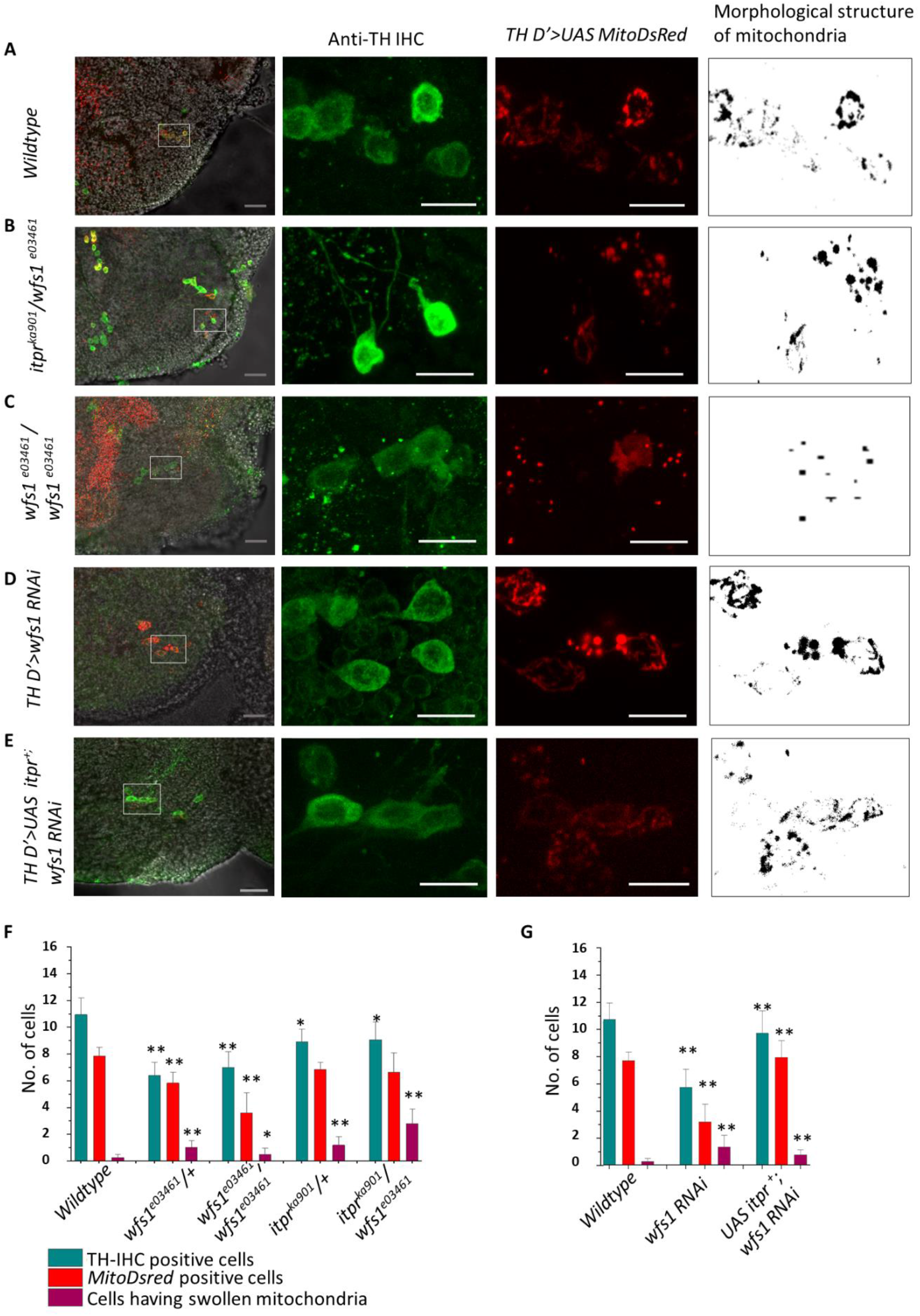
Loss of dopaminergic neurons and altered mitochondrial morphology in *wfs1* and itpr deficient conditions. (A-E) From left to right, representative images of the PPL1 cluster of dopaminergic neurons in a 5 day old fly brain further magnified and visualised by maximum intensity projections of z-stacks of anti-Tyrosine hydroxylase (TH) immunostaining (green) and *TH D’ >MitoDsRed* (Red), in the indicated genotypes. For each genotype the brains and PPL1 cells visualised were as follows: wildtype (N=11, n=117, *itpr* ^*ka901*^*/wfs1*^*e03461*^ transheterozygotes (N=11, n=97), *wfs1*^*e03461*^*/wfs1*^*e03461*^ null mutants (N=10, n=68), *wfs1 RNAi* (N=12, n=68) and (E) *UAS itpr*^*+*^ rescue of *wfs1 RNAi* (N=8, n=77). (F) Graphical representation of proportion of *MitoDsRed*-positive cells in PPL1 neuronal clusters indicates greater loss of mitochondria in the indicated genotypes and (G) Swollen mitochondria in wildtype controls, *itpr* and *wfs1 RNAi* and their corresponding over-expression lines, heterozygotic controls, *wfs1* null mutants (*wfs1*^*e03461*^*/wfs1*^*e03461*^) and *itpr* ^*ka901*^*/wfs1*^*e03461*^ transheterozygotes (N≥8 brains per genotype, n≥ 50 cells per genotype,** p-values <0.05,Student’s t-test and Kruskal-Wallis test).

Next, we observed the mitochondrial morphology of *TH D’* neurons with *wfs1* and *itpr* knockdown. In both cases, the mitochondria appeared swollen (Fig 5D, Supp 4 C). Cells with swollen mitochondria showed anincrease from 0.27+/-1.5 in wildtype PPL1 cells to 1.45+/-0.58 in *wfs1*^*RNAi*^. TH staining indicated lower numbers of TH-stained cells upon knockdown of either wfs1 ((*wfs1*^*RNAi*^) or the IP_3_R (*itpr* ^*RNAi*^; Fig 5E).

As indicated before (Fig 1D), flies with *wfs1*^*RNAi*^ and *itpr*^*RNAi*^ expression in TH D’ cells exhibit comparable flight defects. Both these phenotypes can be rescued by over expression of *UAS itpr*^*+*^ and *UAS wfs1*^*+*^ respectively (Fig 5 E, Supp 4D). To understand if defects in mitochondrial morphology observed in *wfs1 RNAi* can be rescued by overexpression of *itpr*^*+*^, we looked at mitochondrial morphology in the neurons of rescued genotypes. We co-expressed *UAS itpr*^*+*^, *wfs1 RNAi* and *MitodsRed* using *THD’ GAL4* driver. Additionally anti-TH staining was performed on the samples to visualize dopaminergic neurons in these brains. The overexpression of *itpr*^*+*^ restored the mitochondrial morphology and number of TH positive cells in the PPL1 neuronal cluster, caused by knockdown of *wfs1* with *THD’GAL4* (Fig 5E). The rescues indicated a restoration of number of cells with healthy mitochondrial morphology. The DsRed positivity increased from 3.45+/-0.87 in *wfs1*^*RNAi*^ to 7.87+/-0.75in *wfs1*^*RNAi*^ co-expressed with *itpr*^*+*^. The number of cells with swollen mitochondria also decreased from 2.46+/-0.57 in *wfs1*^*RNAi*^ to 0.75+/-0.70 when co-expressed with *itpr* ^*+*^.

Similarly, in flies with knockdown of *itpr* in THD’ neurons, expression of *UAS wfs1*^*+*^ led to a rescue in mitochondrial morphology and other neuronal phenotypes. The cells with mitochondrial DsRed went up from 3.41+/-0.56 to 4.18+/-0.35 when *wfs1*^*+*^ was over expressed. Overall, decrease in dopaminergic neurons and increase in number of deformed or swollen mitochondria in *wfs1 RNAi* and *itpr RNAi* were rescued by overexpression of the other transgene. The null mutants (*wfs1*^*e03461*^*/wfs1*^*e03461*^) on the other hand indicated a decreased level of mitochondria among the TH-IHC stained PPL1 cells (7.63+/-0.81 in wildtype to 3.5+/-0.98 in *wfs1*^*e03461*^*/wfs1*^*e03461*^) in addition to the decrease in neuronal numbers(10.63+/-0.81 in wildtype to 6.8+/-0.77 in *wfs1*^*e03461*^*/wfs1*^*e03461*^) (Fig 5B).

## Discussion

Wolframin or WFS1 is a regulator of Ca^2+^ homeostasis. It interferes with functioning of IP_3_R and SERCA and additionally impacts Endoplasmic reticular stress. In this study, we focus on the interaction between WFS1 and IP_3_R in the context of *Drosophila* flight. We hypothesised that WFS1 is required for maintaining flight bouts in *Drosophila* as was previously reported with IP_3_R. The released Ca^2+^ from Endoplasmic Reticulum through the IP_3_R reaches the cytoplasm or mitochondria where it is required as co-factors for enzymes, for synaptic transmission, muscle contraction, transcription and as second messengers (Berridge *et al*., 2003). An important sink for Ca^2+^ is mitochondria where it acts as co-factor for enzymes for Kreb’s cycle and Oxidative Phosphorylation and for translocases (Rossi *et al*., 2019). Various mitochondrial malfunctions have been reported in the case of neurodegenerative diseases. Dysfunctional mitochondria may trigger or delay mitochondrial turnover, apoptosis, axonal transport and ER damage (Paillusson *et al*., 2016). We hypothesised impaired Ca^2+^ homeostasis due to WFS1 deficiency could be having such an effect on neurons involved in flight bout maintenance.

### WFS1 is involved in flight bout maintenance

Our findings suggest the requirement of WFS1 for flight in *Drosophila* in an IP_3_R dependent manner. *wfs1* null mutants were completely unable to fly from early stages, whereas truncated *wfs1* was able to sustain flight as well as wildtype controls until later stages (20 days). Additionally, the use of *wfs1* knockdown (*wfs1 RNAi*) revealed flight defects comparable to that of IP_3_R knockdown (*itpr RNAi*).

The requirement of WFS1 as an interacting partner of IP_3_R has been established well in other model systems. Our studies demonstrate genetic interactions between the two in *Drosophila* as well. The use of transheterozygotic combination of a *wfs1* mutant allele and *itpr* mutant allele showed a significant decrease in flight duration when compared to the heterozygotic controls. The overexpression analysis using *UAS wfs1*^*+*^ and *UAS itpr*^*+*^ in knockdown and combinations of mutants gave a improvement in flight durations.

### The requirement of WFS1 in drosophila flight is spatially localized majorly to dopaminergic subset of neurons

WFS1 has also been shown to be involved in having an effect on dopamine release system in mouse models. The dopaminergic neurons in vertebrates have known to be involved in modulating locomotion. A classic example is Parkinson’s disease where dopaminergic neurons disintegrate and results in locomotory defects (Stack and Ashburn, 2008). Reports also show dependency of WFS1 on Pink1-Parkin mechanism, where a rescue of wfs1 occurs with the overexpression of Pink1 or Parkin shRNA in Primary rat neuronal cultures (Cagalinec *et al*., 2016). Furthermore, studies done prior to date indicates the requirement of IP_3_R mediated Ca^2+^ homeostasis in dopaminergic neurons is sufficient for flight. Our studies show a similar requirement of WFS1 in dopaminergic subset of neurons. The phenotypes generated by panneuronal *nSybGAL4* was comparable to that generated by dopaminergic *TH GAL4* and subset *TH D’ GAL4*. We also over-expressed *UAS wfs1*^*+*^ and *UAS itpr* ^+^ specifically in dopaminergic *TH D’* neurons and found a rescue of the flight defects caused by deficiency caused by mutants and knock-down.

### WFS1 deficiency impacts Ca^2+^ homeostasis

IP_3_R is specifically involved in Endoplasmic Reticular Ca^2+^ release upon signalling cues. It functions as a channel and also as a component of the Mitochondria Associated Membrane (MAM). Hence, the release of Ca^2+^ and ER-Mitochondrial tethering could be two mechanisms involved in the mechanism of WFS1-IP_3_R interaction, both of which has been demonstrated in rodent systems as well as patient cell cultures.

Carbachol induced Ca^2+^ release experiments result in a decrease in the number of responding cells in *wfs1 RNAi* and *itpr RNAi*. Further, the cells that respond show unregulated Ca^2+^ release as indicated by decreased rate of Ca^2+^ release and fluctuations. Both cytoplasmic and mitochondrial Ca^2+^ responses are decreased upon using the knockdown.

### Mitochondrial health is negatively affected in WFS1 deficient flies

Disruption of Ca^2+^ homeostasis has been shown to result in mitochondrial dysfunction and neuronal loss in various neurodegenerative diseases. The analysis of mitochondrial morphology is the foremost method used as a primary indicator of mitochondrial health. Any mitochondrial functional impairment would result in triggering mechanisms needed for their removal which includes fission-fusion mechanism. Observations of mitochondria indicated a decrease in mitochondrial content and swollen, enlarged mitochondria in the cells deficient in WFS1 and IP_3_R. This abnormally swollen mitochondria were found in transheterozygotes of *itpr* and *wfs1* mutants as well, indicating the interaction is required for maintenance of healthy mitochondria. Additionally, there was a decrease in the number of neurons positive for dopamine.

From the above results we propose that WFS1 mediated Ca^2+^ homeostasis is required for mitochondrial function and thus for neuronal health.

In conclusion, our reverse genetic study of WFS1 suggests that it is required for the functioning of IP_3_R and ER Ca^2+^ release by IP_3_R in *Drosophila*. We demonstrate genetic interactions between IP_3_R and WFS1 through transheterozygotic and over-expression analysis. We propose that interaction between WFS1 and IP_3_R are required for maintaining flight bouts in *Drosophila* and that the requirement is specifically required in the dopaminergic subset of neurons previously reported to be involved in flight circuit functioning. The ER-Ca^2+^ release slows down in these neurons as shown by use of external chemical stimuli. The mitochondrial morphology and content in these neurons within the flight defective genotypes appear perturbed.

## Graphical Abstract

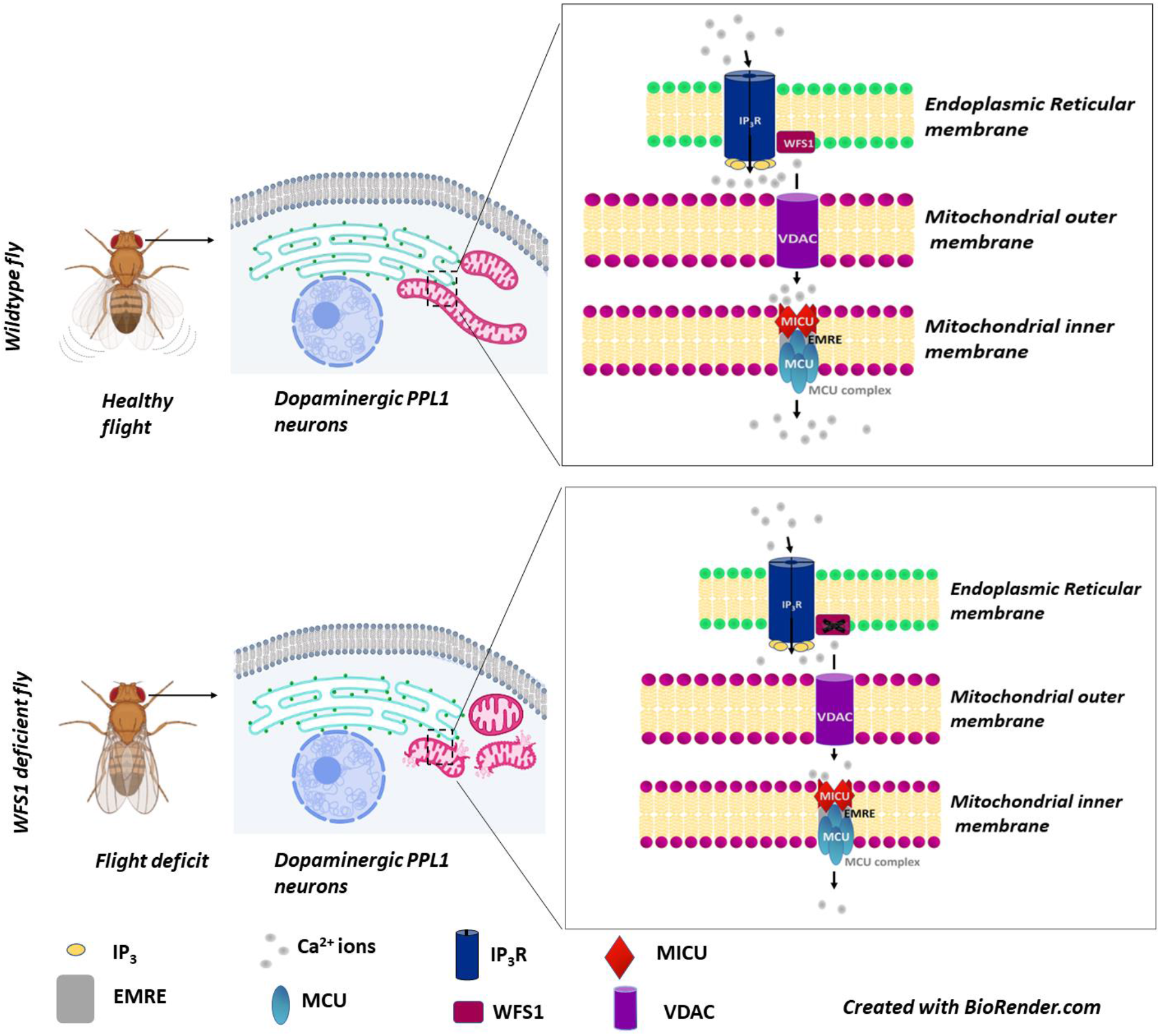

## Legends for supplementary figures

**Supp 1:**
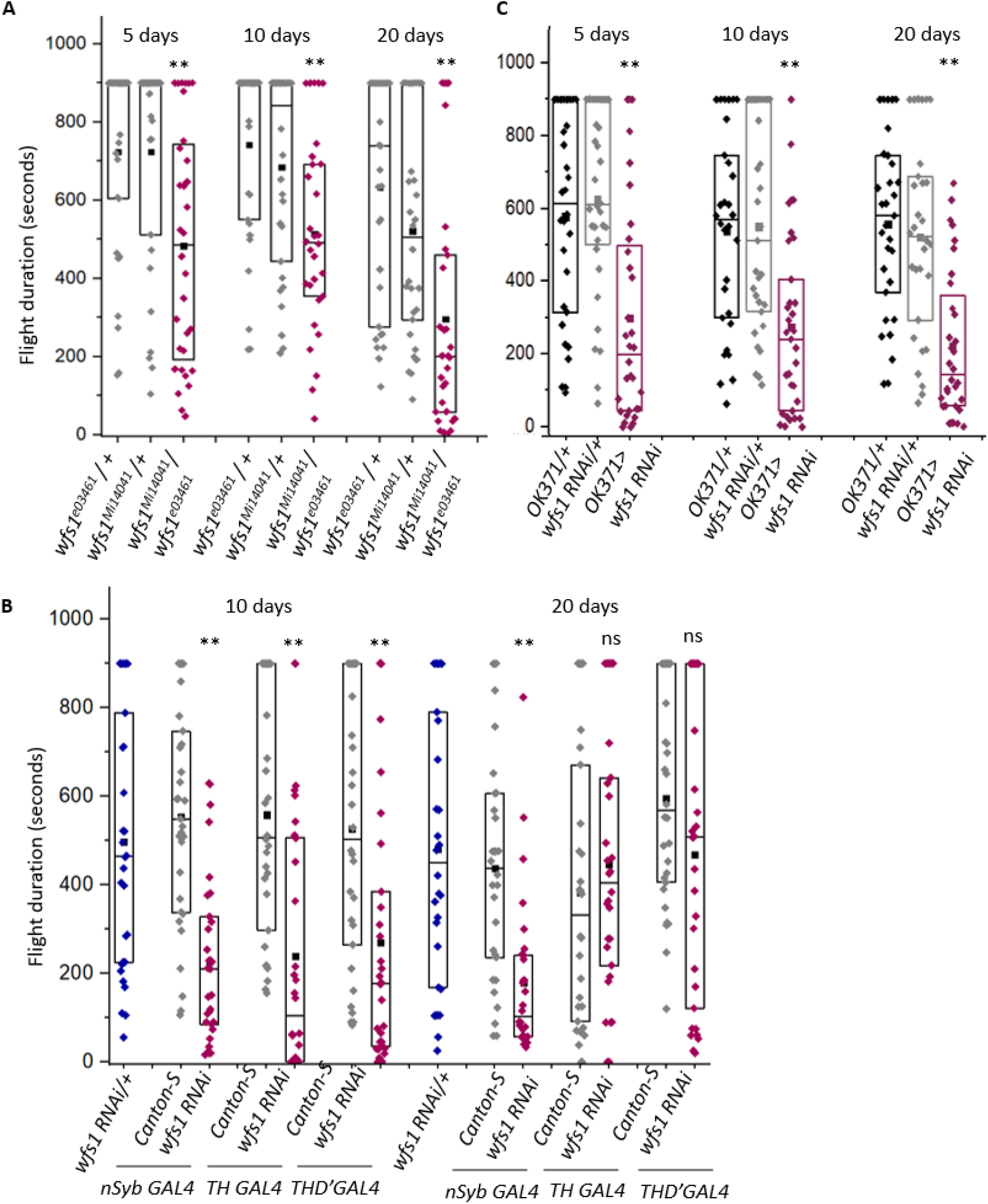
(A) Graphical representation of flight defects in heteroallelic combinations of wfs1 null mutants and wfs1 hypomorphs (B) *GAL4 –UAS RNAi* mediated knockdown of *wfs1* using pan neuronal *GAL4(nSyb), THGAL4* and *TH D’GAL4* (10 days and 20 days old) and (C) glutamatergic *OK371 GAL4*(5, 10 and 20 days old, N ≥30) (** p-values 0.001-0.005, Mann-Whitney test).

**Supp 2:**
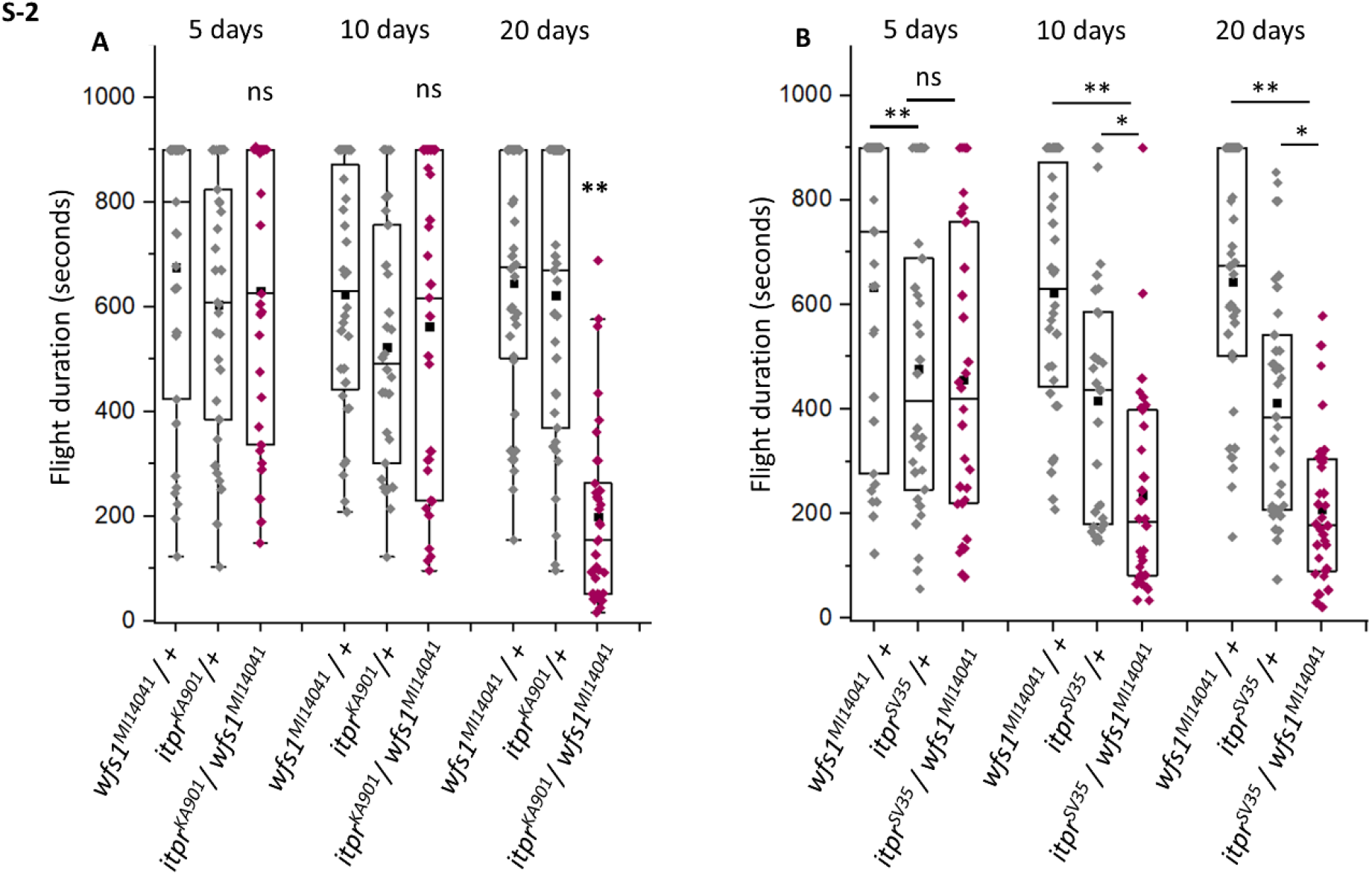
Age dependant flight deficits in *wfs1* ^*MI14041*^, a C-terminal truncated mutant of *wfs1* consisting of 592 aa, in transheterozygotic combination with (A) *itpr* ^*ka901*^, harbouring a point mutation in the Ca^2+^ channel; And (B) *itpr* ^*sv35*^, a truncated mutant of IP_3_R containing 1572 aa. (N ≥30, *** p-values < 0.005, ** p-values 0.001-0.005, Mann-Whitney test).

**Supp 3:**
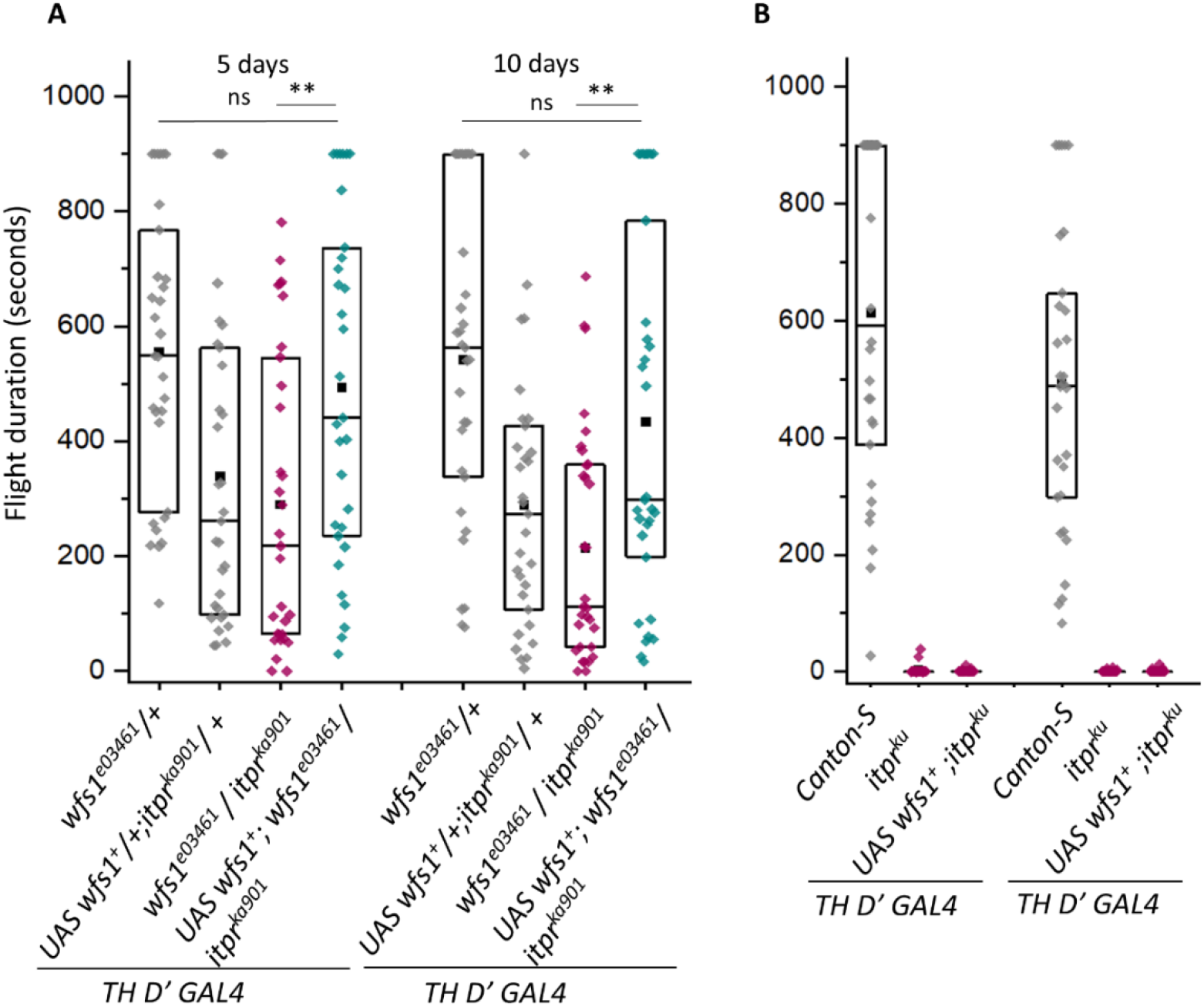
TH D’ *GAL4* mediated overexpression of WFS1 in transheterozygotes and *itpr*^*ku*^ mutants. (A) Rescue of *(wfs1*^*e03461*^/+; *itpr*^*ka901*^/+) by overexpression of WFS1 driven by *TH D’ GAL4* (B) Over expression of WFS1 in *itpr* ^*ku*^ flies(N ≥30, ** p-values 0.001, Mann-Whitney test).

**Supp 4:**
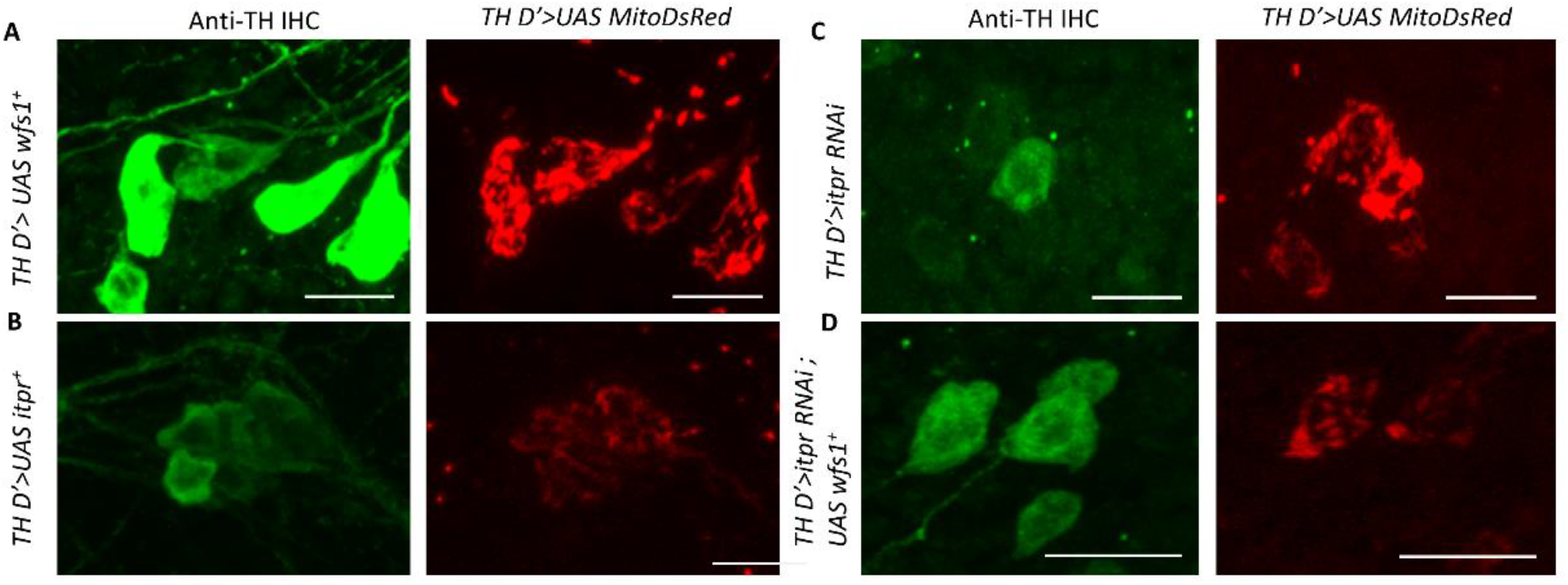
Mitochondrial morphology of PPL1 cluster of neurons i. (A) anti-Tyrosine hydroxylase (TH) immunostaining and visualisation of mitochondria (*TH D’>MitoDsRed*) in 5 day old fly brains of (A) *TH D’>UAS wfs1*^*+*^ over-expression, (B) *TH D’>UAS itpr*^*+*^ over-expression, (C) *itpr RNAi* and (D) *UAS wfs1*^*+*^ rescue of *itpr RNAi*. Number of cells containing (N≥6 brains per genotype, n≥ 50 cells per genotype).

**Supp table -1.**

| Genotype | Wildtype | *wfs1*<sup>RNAi</sup> | *itpr*<sup>RNAi</sup> |
| --- | --- | --- | --- |
| No. of brains used (N) | 5 | 6 | 5 |
| Total no. of cells (n) | 31 | 36 | 29 |
| **JRCaMP counts** |  |  |  |
| Early responders | 6 | 7 | 5 |
| Intermediate responders | 4 | 0 | 1 |
| Late responders | 2 | 3 | 4 |
| Non-responders | 19 | 26 | 19 |
| **MitoGCaMP counts** |  |  |  |
| Early responders | 9 | 5 | 1 |
| Intermediate responders | 7 | 0 | 0 |
| Late responders | 5 | 2 | 10 |
| Non-responders | 10 | 29 | 18 |

## References

1. Amr, S., Heisey, C., Zhang, M., Xia, X.J., Shows, K.H., Ajlouni, K., Pandya, A., Satin, L.S., El-Shanti, H. and Shiang, R., 2007. A homozygous mutation in a novel zinc-finger protein, ERIS, is responsible for Wolfram syndrome The American Journal of Human Genetics, 81(4): pp.673–683.

2. Angebault, C., Fauconnier, J., Patergnani, S., Rieusset, J., Danese, A., Affortit, C.A., Jagodzinska, J., Mégy, C., Quiles, M., Cazevieille, C. and Korchagina, J., 2018. ER-mitochondria cross-talk is regulated by the Ca2+ sensor NCS1 and is impaired in Wolfram syndrome. Science signaling, 11(553): p.eaaq1380.

3. Banerjee, S., Lee, J., Venkatesh, K., Wu, C.F. and Hasan, G., 2004. Loss of flight and associated neuronal rhythmicity in inositol 1, 4, 5-trisphosphate receptor mutants of Drosophila. Journal of Neuroscience, 24(36), pp.7869–7878.

4. Barazzuol, L., Giamogante, F., Brini, M. and Calì, T., 2020. PINK1/parkin mediated mitophagy, Ca2+ signalling, and ER–mitochondria contacts in Parkinson’s disease. International journal of molecular sciences, 21(5), p.1772.

5. Berridge, M.J., 2003. Cardiac calcium signalling. Biochemical Society Transactions, 31(5), pp.930–933.

6. Cagalinec, M., Liiv, M., Hodurova, Z., Hickey, M.A., Vaarmann, A., Mandel, M., Zeb, A., Choubey, V., Kuum, M., Safiulina, D. and Vasar, E., 2016. Role of mitochondrial dynamics in neuronal development: mechanism for Wolfram syndrome. PLoS biology, 14(7), p.e1002511.

7. Cremers, C.W., Wijdeveld, P.G. and Pinckers, A.J., 1977. Juvenile diabetes mellitus, optic atrophy, hearing loss, diabetes insipidus, atonia of the urinary tract and bladder, and other abnormalities (Wolfram syndrome). A review of 88 cases from the literature with personal observations on 3 new patients. Acta paediatrica Scandinavica. Supplement, (264), pp.1–16.

8. Friggi-Grelin, F., Coulom, H., Meller, M., Gomez, D., Hirsh, J. and Birman, S., 2003. Targeted gene expression in Drosophila dopaminergic cells using regulatory sequences from tyrosine hydroxylase. Journal of neurobiology, 54(4), pp.618–627.

9. Hom, J., Yu, T., Yoon, Y., Porter, G. and Sheu, S.S., 2010. Regulation of mitochondrial fission by intracellular Ca2+ in rat ventricular myocytes. Biochimica et Biophysica Acta (BBA)-Bioenergetics, 1797(6-7), pp.913–921.

10. Inoue, H., Tanizawa, Y., Wasson, J., Behn, P., Kalidas, K., Bernal-Mizrachi, E., Mueckler, M., Marshall, H., Donis-Keller, H., Crock, P. and Rogers, D., 1998. A gene encoding a transmembrane protein is mutated in patients with diabetes mellitus and optic atrophy (Wolfram syndrome). Nature genetics, 20(2), pp.143–148.

11. Joshi, R., Venkatesh, K., Srinivas, R., Nair, S. and Hasan, G., 2004. Genetic dissection of itpr gene function reveals a vital requirement in aminergic cells of Drosophila larvae. Genetics, 166(1), pp.225–236.

12. Kakiuchi, C., Ishiwata, M., Hayashi, A. and Kato, T., 2006. XBP1 induces WFS1 through an endoplasmic reticulum stress response element-like motif in SH-SY5Y cells. Journal of neurochemistry, 97(2), pp.545–555.

13. Lezi, E. and Swerdlow, R.H., 2012. Mitochondria in neurodegeneration. Advances in Mitochondrial Medicine, pp.269–286.

14. Liu, Q., Liu, S., Kodama, L., Driscoll, M.R. and Wu, M.N., 2012. Two dopaminergic neurons signal to the dorsal fan-shaped body to promote wakefulness in Drosophila. Current Biology, 22(22), pp.2114–2123.

15. Manjila, S.B. and Hasan, G., 2018. Flight and climbing assay for assessing motor functions in Drosophila. Bio-protocol, 8(5), pp.e2742–e2742.

16. Mao, Z. and Davis, R.L., 2009. Eight different types of dopaminergic neurons innervate the Drosophila mushroom body neuropil: anatomical and physiological heterogeneity. Frontiers in neural circuits, 3, p.5.

17. Matsunaga, K., Tanabe, K., Inoue, H., Okuya, S., Ohta, Y., Akiyama, M., Taguchi, A., Kora, Y., Okayama, N., Yamada, Y. and Wada, Y., 2014. Wolfram syndrome in the Japanese population; molecular analysis of WFS1 gene and characterization of clinical features. PloS one, 9(9), p.e106906.

18. Matto, V., Terasmaa, A., Vasar, E. and Kõks, S., 2011. Impaired striatal dopamine output of homozygous Wfs1 mutant mice in response to [K+] challenge. Journal of physiology and biochemistry, 67(1), pp.53–60.

19. Millar, C., Poyer, J.F., Gabelt, B.T. and Kaufman, P.L., 1995. Endothelin subtypes: effect on isolated rhesus monkey ciliary muscle. Journal of Pharmacology and Experimental Therapeutics, 275(3), pp.1143–1147.

20. Odisho, T., Zhang, L. and Volchuk, A., 2015. ATF6β regulates the Wfs1 gene and has a cell survival role in the ER stress response in pancreatic β-cells. Experimental cell research, 330(1), pp.111–122.

21. Osman, A.A., Saito, M., Makepeace, C., Permutt, M.A., Schlesinger, P. and Mueckler, M., 2003. Wolframin expression induces novel ion channel activity in endoplasmic reticulum membranes and increases intracellular calcium. Journal of Biological Chemistry, 278(52), pp.52755–52762.

22. Paillusson, S., Stoica, R., Gomez-Suaga, P., Lau, D.H., Mueller, S., Miller, T. and Miller, C.C., 2016. There’s something wrong with my MAM; the ER– mitochondria axis and neurodegenerative diseases. Trends in neurosciences, 39(3), pp.146–157.

23. Pathak, T., Agrawal, T., Richhariya, S., Sadaf, S. and Hasan, G., 2015. Store-operated calcium entry through Orai is required for transcriptional maturation of the flight circuit in Drosophila. Journal of Neuroscience, 35(40), pp.13784–13799.

24. Rangaraju, V., Lewis, T.L., Hirabayashi, Y., Bergami, M., Motori, E., Cartoni, R., Kwon, S.K. and Courchet, J., 2019. Pleiotropic mitochondria: the influence of mitochondria on neuronal development and disease. Journal of Neuroscience, 39(42), pp.8200–8208.

25. Ravi, P., Trivedi, D. and Hasan, G., 2018. FMRFa receptor stimulated Ca2+ signals alter the activity of flight modulating central dopaminergic neurons in Drosophila melanogaster. PLoS genetics, 14(8), p.e1007459.

26. Rohayem, J., Ehlers, C., Wiedemann, B., Holl, R., Oexle, K., Kordonouri, O., Salzano, G., Meissner, T., Burger, W., Schober, E. and Huebner, A., 2011. Diabetes and neurodegeneration in Wolfram syndrome: a multicenter study of phenotype and genotype. Diabetes care, 34(7), pp.1503–1510.

27. Rossi, A., Pizzo, P. and Filadi, R., 2019. Calcium, mitochondria and cell metabolism: A functional triangle in bioenergetics. Biochimica et Biophysica Acta (BBA)-Molecular Cell Research, 1866(7), pp.1068–1078.

28. Rouzier, C., Moore, D., Delorme, C., Lacas-Gervais, S., Ait-El-Mkadem, S., Fragaki, K., Burté, F., Serre, V., Bannwarth, S., Chaussenot, A. and Catala, M., 2017. A novel CISD2 mutation associated with a classical Wolfram syndrome phenotype alters Ca2+ homeostasis and ER-mitochondria interactions. Human molecular genetics, 26(9), pp.1599–1611.

29. Sakakibara, Y., Sekiya, M., Fujisaki, N., Quan, X. and Iijima, K.M., 2018. Knockdown of wfs1, a fly homolog of Wolfram syndrome 1, in the nervous system increases susceptibility to age-and stress-induced neuronal dysfunction and degeneration in Drosophila. PLoS genetics, 14(1), p.e1007196.

30. Scolding, N.J., Kellar-Wood, H.F., Shaw, C., Shneerson, J.M. and Antount, N., 1996. Wolfram syndrome: hereditary diabetes mellitus with brainstem and optic atrophy. Annals of Neurology: Official Journal of the American Neurological Association and the Child Neurology Society, 39(3), pp.352–360.

31. Sharma, A. and Hasan, G., 2020. Modulation of flight and feeding behaviours requires presynaptic IP3Rs in dopaminergic neurons. Elife, 9, p.e62297.

32. Srikanth, S., Banerjee, S. and Hasan, G., 2006. Ectopic expression of a Drosophila InsP3R channel mutant has dominant-negative effects in vivo. Cell calcium, 39(2), pp.187–196.

33. Stack, E. and Ashburn, A., 2008. Dysfunctional turning in Parkinson’s disease. Disability and rehabilitation, 30(16), pp.1222–1229.

34. Strom, T.M., Hörtnagel, K., Hofmann, S., Gekeler, F., Scharfe, C., Rabl, W., Gerbitz, K.D. and Meitinger, T., 1998. Diabetes insipidus, diabetes mellitus, optic atrophy and deafness (DIDMOAD) caused by mutations in a novel gene (wolframin) coding for a predicted transmembrane protein. Human molecular genetics, 7(13), pp.2021–2028.

35. Tekko, T., Lakspere, T., Allikalt, A., End, J., Kolvart, K.R., Jagomäe, T., Terasmaa, A., Philips, M.A., Visnapuu, T., Väärtnõu, F. and Gilbert, S.F., 2017. Wfs1 is expressed in dopaminoceptive regions of the amniote brain and modulates levels of D1-like receptors. PloS one, 12(3), p.e0172825.

36. Tranebjaerg, L., Barrett, T. and Rendtorff, N.D., 2009. WFS1–related disorders, Pagon RA, Bird TC, Dolan CR, Stephens K, eds. Gene Reviews (internet).

37. Valente, A. J., Maddalena, L. A., Robb, E. L., Moradi, F., & Stuart, J. A. (2017). A simple ImageJ macro tool for analyzing mitochondrial network morphology in mammalian cell culture. Acta histochemica, 119(3), 315–326.

38. Visnapuu, T., Plaas, M., Reimets, R., Raud, S., Terasmaa, A., Kõks, S., Sütt, S., Luuk, H., Hundahl, C.A., Eskla, K.L. and Altpere, A., 2013. Evidence for impaired function of dopaminergic system in Wfs1-deficient mice. Behavioural brain research, 244, pp.90–99.

